## Supplementary Material for "Body size as a metric for the affordable world"

**This PDF file includes:**

Figs. S1 to S4
Tables S1 to S6

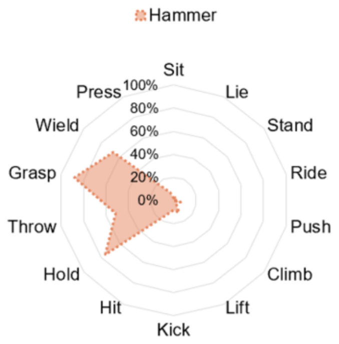


**Fig. S1.** Two exemplar objects with different affordance profiles. The values represented the percentage of participants who agreed on a certain action being afforded by an object.

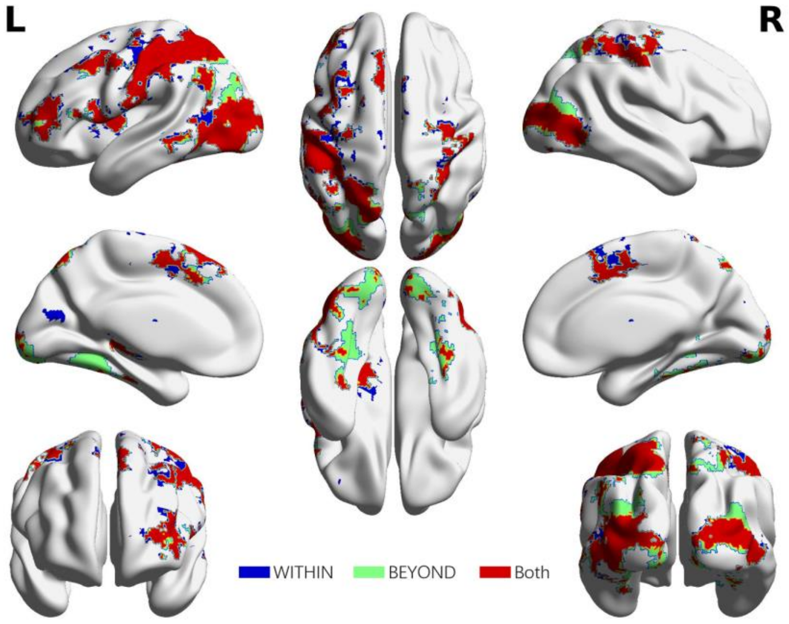


**Fig. S2.** Brain areas showing significantly greater neural activation for objects within body size and beyond body size versus baseline.

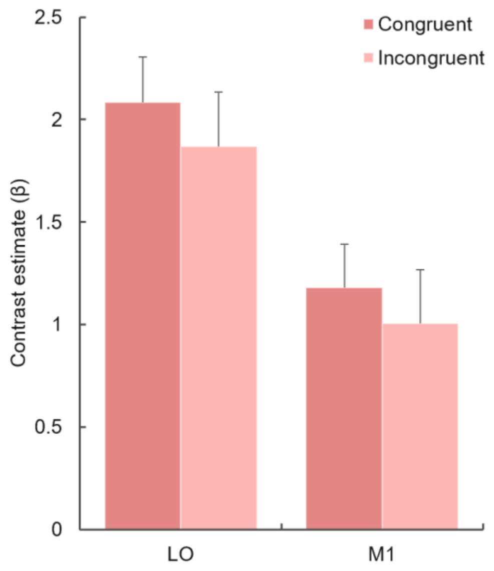


**Fig. S3.** The activation in LO and M1 in response to objects **within** body size in the congruent and incongruent conditions, respectively. The bars refer to the contrast estimates of each condition versus baseline. The stars indicate whether the contrast was significant. *$p$<.05, **$p$<.01, ***$p$<.001, otherwise not significant. Error bars represent the standard error (SE).

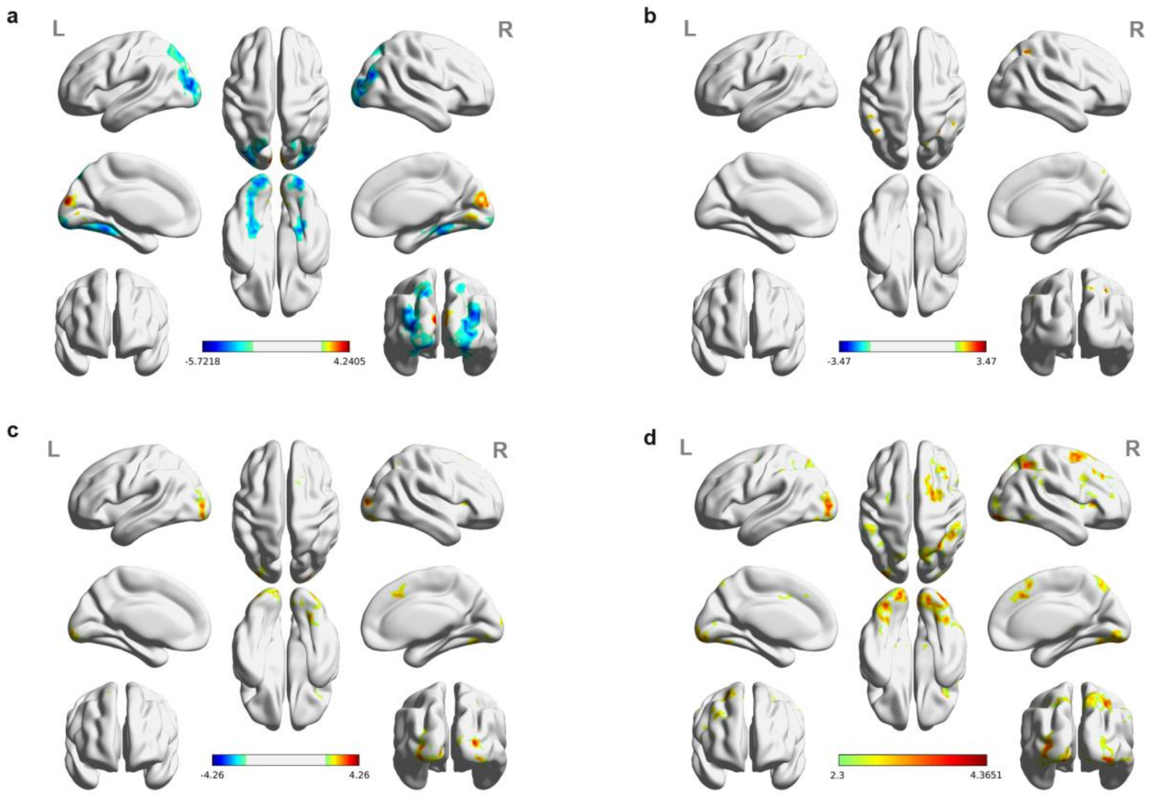


**Fig. S4.** Significant brain activations of different contrasts in the whole-brain level analysis. **a,** the effect of object type, positive values (warm color) indicated higher activation for objects within than objects beyond and negative values (cold color) indicated the opposite. **b,** the effect of congruency, positive values indicated higher activation in congruent than incongruent condition. **c,** the effect of interaction between object type and congruency, positive values indicated the larger congruency effect for objects within than beyond. **d,** the congruency effect for objects within. All contrasts were corrected with cluster-level correction at $p < .05$. The detailed cluster-level results for each contrast map can be found in Supplementary Table S2 to S5.

**Table S1**. Cortical regions showing object-selective activation for whole-brain conjunction analysis (R = right hemisphere, L = left hemisphere; Z > 2.3, *p* = 0.05, cluster corrected)

| Cluster | Region | Number of voxels in region | MNI coordinates | | | Peak Z value |
|---|---|---|---|---|---|---|
| | | | x | y | z | |
| 1 | L Postcentral Gyrus | 1804 | -54 | -26 | 50 | 5.41 |
| | L Middle Occipital Gyrus | 1392 | | | | |
| | L Inferior Parietal Lobule | 1321 | | | | |
| | L Superior Parietal Lobule | 1284 | | | | |
| | L Supplementary Motor Area | 851 | | | | |
| | L Precentral Gyrus | 749 | | | | |
| | L Inferior Occipital Gyrus | 409 | | | | |
| | L Middle Temporal Gyrus | 402 | | | | |
| | L Fusiform Gyrus | 390 | | | | |
| | L Precuneus | 320 | | | | |
| | L Inferior Temporal Gyrus | 303 | | | | |
| 2 | R Middle Occipital Gyrus | 594 | 28 | -90 | 12 | 4.55 |
| | R Middle Temporal Gyrus | 335 | | | | |
| | R Fusiform Gyrus | 321 | | | | |
| | R Inferior Temporal Gyrus | 209 | | | | |
| | R Inferior Occipital Gyrus | 145 | | | | |
| 3 | R Postcentral Gyrus | 896 | 30 | -58 | 58 | 3.90 |
| | R Precentral Gyrus | 577 | | | | |
| | R Superior Parietal Lobule | 427 | | | | |
| 4 | L Rolandic Operculum | 331 | -50 | 8 | 4 | 3.97 |
| | L Inferior Frontal Gyrus, opercular part | 268 | | | | |
| 5 | L Middle Frontal Gyrus | 520 | -28 | 46 | 8 | 3.68 |
| | L Inferior Frontal Gyrus, triangular part | 245 | | | | |

**Table S2**. Cortical regions reaching significance in the contrasts of (A) objects within versus object beyond and (B) objects beyond versus objects within, whole-brain analysis (R = right hemisphere, L = left hemisphere; Z > 2.3, *p* = 0.05, cluster corrected)

| **Cluster** | **Region** | **Number of voxels in region** | **MNI coordinates** | | | **Peak Z value** |
|---|---|---|---|---|---|---|
| | | | **x** | **y** | **z** | |
| Objects within > Objects beyond | | | | | | |
| 1 | L Cuneus | 363 | 10 | -90 | 20 | 4.24 |
| | R Cuneus | 309 | | | | |
| | L Lingual Gyrus | 176 | | | | |
| | R Lingual Gyrus | 293 | | | | |
| Objects beyond > Objects within | | | | | | |
| 1 | L Middle Occipital Gyrus | 1831 | -16 | -92 | -8 | 5.72 |
| | L Fusiform Gyrus | 994 | | | | |
| | L Superior Parietal Lobule | 584 | | | | |
| | L Lingual Gyrus | 504 | | | | |
| | L Inferior Occipital Gyrus | 304 | | | | |
| | L Superior Occipital Gyrus | 211 | | | | |
| | L Parahippocampal Gyrus | 210 | | | | |
| | L Precuneus | 205 | | | | |
| 2 | R Middle Occipital Gyrus | 1358 | 20 | -86 | -2 | 5.40 |
| | R Lingual Gyrus | 340 | | | | |
| | R Superior Parietal Lobule | 328 | | | | |
| | R Superior Occipital Gyrus | 320 | | | | |
| | R Inferior Occipital Gyrus | 276 | | | | |
| 3 | R Fusiform Gyrus | 483 | 34 | -38 | -16 | 4.73 |
| | R Parahippocampal Gyrus | 316 | | | | |

**Table S3**. Cortical regions reaching significance in contrasts of (A) congruent versus incongruent and (B) incongruent versus congruent, whole-brain analysis (R = right hemisphere, L = left hemisphere; Z > 2.3, *p* = 0.05, cluster corrected)

| Cluster | Region | Number of voxels in region | MNI coordinates | | | Peak Z value |
|---|---|---|---|---|---|---|
| | | | x | y | z | |
| Congruent > Incongruent | | | | | | |
| 1 | L Inferior Parietal Lobule | 322 | -44 | -50 | 64 | 3.47 |
| 2 | R Superior Parietal Lobule | 339 | 36 | -66 | 52 | 3.31 |
| | R Inferior Parietal Lobule | 165 | | | | |
| Incongruent > Congruent | | | | | | |
| - | *No significant cluster* | - | - | - | - | - |

**Table S4**. Cortical regions showing significant interaction between object type and congruency, whole-brain analysis (OW = Objects within, OB = Objects beyond; R = right hemisphere, L = left hemisphere; Z > 2.3, *p* = 0.05, cluster corrected)

| Cluster | Region | Number of voxels in region | MNI coordinates | | | Peak Z value |
|---|---|---|---|---|---|---|
| | | | x | y | z | |
| (OW_Congruent – OW_Incongruent ) > (OB_Congruent – OB_Incongruent ) | | | | | | |
| 1 | L Middle Occipital Gyrus | 831 | 22 | -94 | 10 | 4.25 |
| | R Middle Occipital Gyrus | 187 | | | | |
| | L Fusiform Gyrus | 113 | | | | |
| | R Fusiform Gyrus | 376 | | | | |
| | L Inferior Occipital Gyrus | 293 | | | | |
| | R Inferior Occipital Gyrus | 276 | | | | |
| | L Lingual Gyrus | 215 | | | | |
| | R Lingual Gyrus | 345 | | | | |
| | L Superior Occipital Gyrus | 123 | | | | |
| 2 | R Supplementary Motor Area | 383 | 14 | 14 | 60 | 3.39 |
| 3 | R Superior Parietal Lobule | 191 | 36 | -62 | 56 | 3.18 |
| | R Inferior Parietal Lobule | 114 | | | | |
| 4 | R Insula | 175 | 32 | 18 | 8 | 3.41 |
| (OB_Congruent – OB_Incongruent ) > (OW_Congruent – OW_Incongruent ) | | | | | | |
| - | *No significant cluster* | - | - | - | - | - |

**Table S5**. Cortical regions showing significant congruency effect (congruent versus incongruent) for objects within, whole-brain analysis (R = right hemisphere, L = left hemisphere; Z > 2.3, *p* = 0.05, cluster corrected)

| Cluster | Region | Number of voxels in region | MNI coordinates | | | Peak Z value |
|---|---|---|---|---|---|---|
| | | | x | y | z | |
| 1 | L Middle Occipital Gyrus | 820 | -14 | -96 | -2 | 4.29 |
| | R Middle Occipital Gyrus | 214 | | | | |
| | L Inferior Occipital Gyrus | 367 | | | | |
| | R Inferior Occipital Gyrus | 487 | | | | |
| | L Lingual Gyrus | 329 | | | | |
| | R Lingual Gyrus | 484 | | | | |
| | L Fusiform Gyrus | 341 | | | | |
| | R Fusiform Gyrus | 463 | | | | |
| | R Inferior Temporal Gyrus | 280 | | | | |
| 2 | R Superior Parietal Lobule | 891 | 32 | -58 | 50 | 3.80 |
| | R Inferior Parietal Lobule | 619 | | | | |
| | R Supramarginal Gyrus | 300 | | | | |
| | R Precuneus | 294 | | | | |
| | L Superior Parietal Lobule | 210 | | | | |
| 3 | L Inferior Parietal Lobule | 668 | -48 | -46 | 60 | 3.82 |
| 4 | R Superior Frontal Gyrus | 501 | 16 | 12 | 60 | 3.57 |
| | R Supplementary Motor Area | 200 | | | | |
| 5 | R Middle Frontal Gyrus | 398 | 40 | 42 | 34 | 3.44 |
| 6 | R Inferior Frontal Gyrus, opercular part | 384 | 50 | 14 | 38 | 3.33 |

**Table S6**. The full list of inanimate objects used in the behavioral study, with the corresponding size rank noted according to Konkle and Oliva (2011).

| Object | Diagonal Size (cm) | Size Rank |
|---|---|---|
| airplane | 7618 | 8 |
| apple | 14 | 2 |
| ball | 40 | 3 |
| bed | 252 | 6 |
| bike | 181 | 5 |
| bottle | 35 | 3 |
| brick | 25 | 3 |
| chair | 115 | 5 |
| eggplant | 31 | 3 |
| hammer | 25 | 3 |
| kettle | 30 | 3 |
| ladder | 213 | 6 |
| laptop | 44 | 4 |
| phone | 17 | 2 |
| piano | 184 | 6 |
| pipa | 107 | 4 |
| plate | 20 | 2 |
| potted plant | 24 | 3 |
| shoe | 32 | 3 |
| skateboard | 83 | 4 |
| sweater | 99 | 4 |
| tent | 500 | 7 |
| tree | 3016 | 8 |
| umbrella | 104 | 4 |
